# Computation on demand: Action-specific representations of visual task features arise during distinct movement phases

**DOI:** 10.1101/2023.11.27.568674

**Authors:** Nina Lee, Lin Lawrence Guo, Adrian Nestor, Matthias Niemeier

## Abstract

It is commonly held that computations of goal-directed behaviour are governed by conjunctive neural representations of the task features. However, support for this view comes from paradigms with arbitrary combinations of task features and task affordances that require representations in working memory. Therefore, in the present study we used a task that is well-rehearsed with task features that afford minimal working memory representations to investigate the temporal evolution of feature representations and their potential integration in the brain. Specifically, we recorded electroencephalography data from human participants while they first viewed and then grasped objects or touched them with a knuckle. Objects had different shapes and were made of heavy or light materials with shape and weight being features relevant for grasping but not for knuckling. Using multivariate analysis, we found that representations of object shape were similar for grasping and knuckling. However, only for grasping did early shape representations reactivate at later phases of grasp planning, suggesting that sensorimotor control signals feed back to early visual cortex. Grasp-specific representations of material/weight only arose during grasp execution after object contact during the load phase. A trend for integrated representations of shape and material also became grasp-specific but only briefly during movement onset. These results argue against the view that goal-directed actions inevitably join all features of a task into a sustained and unified neural representation. Instead, our results suggest that the brain generates action-specific representations of relevant features as required for the different subcomponent of its action computations.

**Significance statement:** The idea that all the features of a task are integrated into a joint representation or event file is widely supported but importantly based on paradigms with arbitrary stimulus-response combinations. Our study is the first to investigate grasping using electroencephalography to search for the neural basis of feature integration in such a daily-life task with overlearned stimulus-response mappings. Contrary to the notion of event files we find limited evidence for integrated representations. Instead, we find that task-relevant features form representations at specific phases of the action. Our results show that integrated representations do not occur universally for any kind of goal-directed behaviour but in a manner of computation on demand.

## Introduction

To interact with the world around us our brain computes action intentions, i.e., cognitive signals (e.g., Andersen & Cui, 2009; Cui, 2014) that modulate our sensorimotor transformations to extract sensory information as needed to produce the desired behaviour (e.g., Bekkering & Neggers, 2002; Craighero et al.,1999; Fagioli et al., 2007; Shin et al., 2010). An important model of goal-directed behaviour is grasping (Grafton, 2010) that has been shown to bias visual input such that attention splits into multiple foci directed to those object parts where the fingers of the grasping hand will touch the object (Brouwer et al., 2009; Schiegg et al., 2003), arguably improving focal processing in visual areas (Baldauf & Deubel, 2009; Baldauf et al., 2006). Indeed, areas in striate and extra-striate cortex carry retinotopically specific information about the impending action (Gallivan et al., 2019). Thus, the intention to grasp seems to be associated with sensorimotor feedback signals that reactivate early visual areas.

Consistent with the idea of reactivation, a recent multivariate EEG study found that participants formed early neural representations of the shape of a grasp target 100-200 ms after an object appeared and, crucially, that the same representations re-occurred around 400-700 ms (Guo et al., 2019). This could indicate that the intention to grasp sets off a detailed visual analysis of the object to extract information about how to guide the grasping fingers to the object.

As another grasp-relevant feature, visual cues about the weight of an object are known to influence load and grip forces during grasping (Baugh et al., 2012; Buckingham et al., 2009; Johansson & Flanagan, 2009). Also, activation of occipitotemporal cortex together with motor and premotor areas has been shown to carry signals about the weight of objects while participants plan to grasp objects (Gallivan et al., 2014).

However, object features such as weight and shape, though grasp-relevant, have yet to be shown to trigger neural representations that are specific to grasping. Alternatively, these representations might form in an automatic manner irrespective of what the actor intends to do. That is, previous studies observed shape and weight representations while participants were planning to grasp but did not compare to scenarios where participants intended to perform different actions (Gallivan et al., 2014; Guo et al., 2019).

What is more, it is unclear whether grasp-specific representations of weight and shape are then integrated into one unified grasp programs. A rich literature has found that various object and task features are joined into “event files” or conjunctive representations (e.g., Hommel, 2019; Kikuomoto & Mayr, 2020; Frings et al., 2020; Hommel, 2022) for the control of goal-directed actions. That said, most of these paradigms tested actions that were based on relatively arbitrary combinations of task features – in contrast to grasping where the use of task features often is overlearned and where object features might be processed selectively (Ganel & Goodale, 2003), potentially eliciting separate representations during different phases of action control (Johansson & Flanagan, 2009).

Thus, the aim of our study was to investigate (a) whether human participants form action-specific representations of task features during different phases of action control, and (b) whether these representations integrate multiple task features. Specifically, we had participants grasp to lift or reach for to knuckle objects that had different shapes and that were made from materials with different densities, i.e., grasp-relevant (but not reach-relevant) features that might elicit grasp-specific representations. Furthermore, we used multivariate analysis of electroencephalography (EEG) recordings because of their high sensitivity for representations of actions at the level of milliseconds (Guo et al., 2019; Sburlea et al., 2021; for similar approach using magnetoencephalography: Turella et al., 2016) and their capability of identifying integrated representations (e.g., Guo et al., 2021; Kikumoto & Mayr, 2020).

## Methods

### Participants

Sixteen undergraduate and graduate students (12 females; median age: 21.5; range: 18-38) from the University of Toronto Scarborough gave their written and informed consent to participate in the experiment and were compensated $15/h for their time. All participants were right-handed (Oldfield, 1971) and had normal or corrected-to-normal vision. All procedures were approved by the Human Participants Review Sub-Committee of the University of Toronto.

### Stimuli

For the experiment we used four simple objects that all were sized to be comfortably graspable along their grasp axis (always 6 cm) but differed noticeably in shape and/or weight, two features that are relevant for grasping objects but not for reaching for objects and touching them with a knuckle. That is, objects had the shapes of “pillows” or “flowers” (Fig. 1A) with either concave (72 deg. segments of circles with a radius of 7.5 cm, i.e., curvature = 13.3/m) or convex sides (180 deg. segments of circles with a radius of 1.5 cm, curvature = 66.7/m). That is, the difference in surface curvature was several orders of magnitude larger than perceptual thresholds and known to yield strongly different grasp performance (e.g., Jenmalm et al., 1998). Furthermore, all shapes were chosen to have two identical symmetry axes to minimize any inherent preferences about grasp orientations.

**Figure 1:**
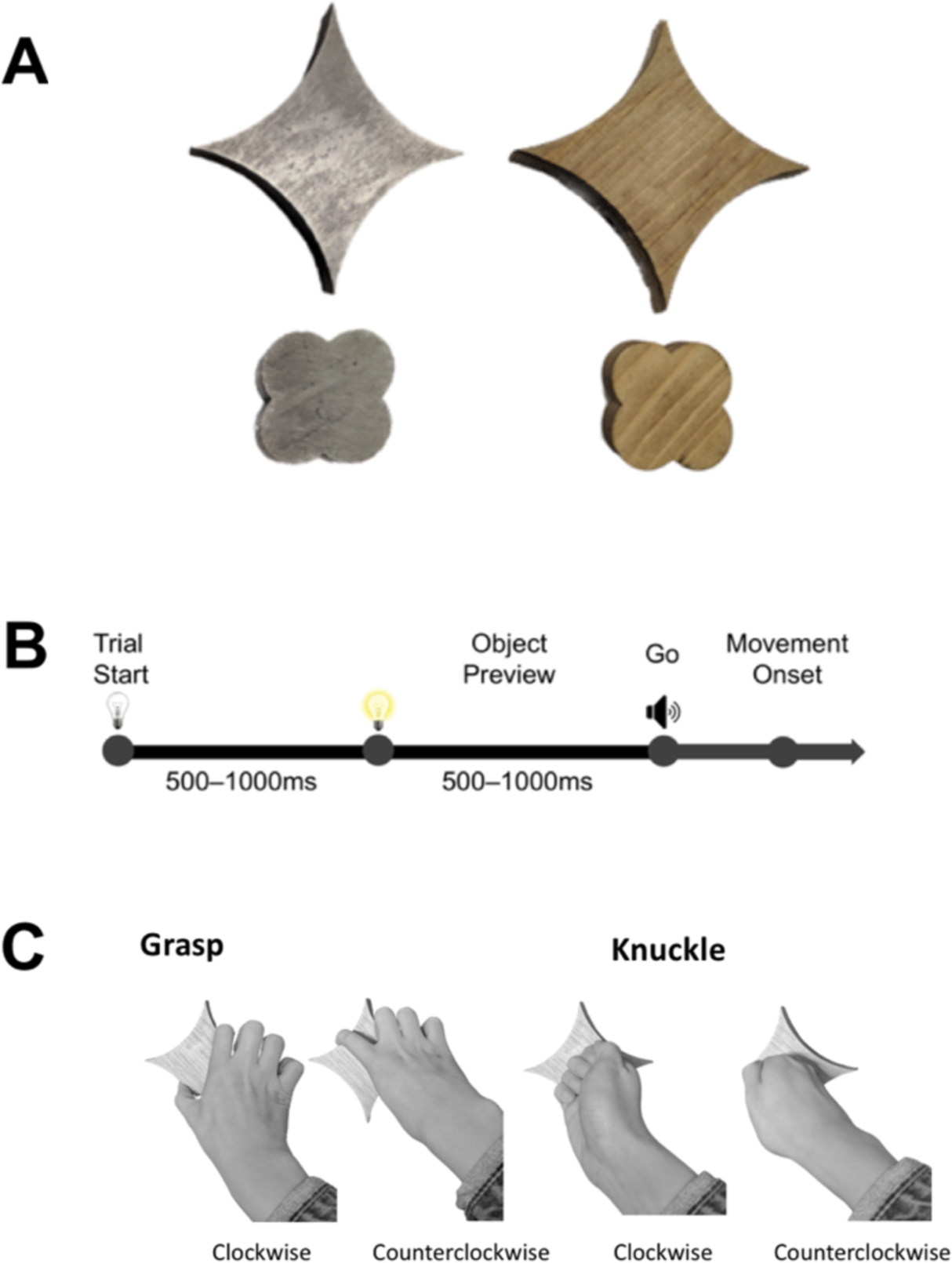
Experimental methods. A. Objects used in the experiment. B. Timeline of a trial. C. Action conditions.

All shapes were either cut from 2 cm thick blocks of wood or steel (machining tolerances of 0.0254 cm) such that on average steel objects were 15.9 heavier than wood objects, a difference that is several orders of magnitude larger than perceptual thresholds of weight (Ellis & Lederman, 1999). (Steel pillow and flower weighed 643 g and 339 g, and were 15.7 and 16.1 times heavier than the wood pillow, 41 g, and the wood flower, 21 g, respectively, due to natural variations in wood density; also, pillows were 1.9 times heavier than flowers due to their larger volume; this difference was noticeable, though much smaller than the material-dependent difference, but was unavoidable given that we aimed for identical grip sizes together with large difference in surface curvature). Furthermore, wood and steel materials were chosen as they intuitively relayed greatly contrasting weight information and were perceptually clearly different in colour and texture.

### Procedure and Apparatus

During the experiment participants sat in a dark room with the experimenter on the left side. To minimize the pre-trial information inadvertently relayed by the experimenter setting up each trial, participants wore earplugs and looked at an opaque shutter glass screen (Smart Class Technology). At the beginning of each trial, participants placed their right index finger and thumb on a button box which obscured a beam of infrared light, signalling the presence of their hand. For the purpose of preparing each trial, the experimenter viewed instructions on a monitor on the side and turned on a set of LEDs to mount objects on a platform in an otherwise dark grasp space. There was a square-shaped hole in the centre of the platform to which objects with square pegs were attached, such that objects were always placed in the same position and orientation, 43 cm away from the participant. Further, the platform was slanted to ensure the surface of the object would be tilted toward the participant’s line of sight.

Upon setting up the object, the experimenter pressed a key that switched off the LEDs and in darkness, set the shutter glass screen to transparent. Five hundred to 1000 ms later, the LED lights turned on, illuminating the object for the participant to see for a “Preview” duration of 500-1000 ms before a “Go” beep signalled participants to make their movement (Fig. 1B). As participants lifted their hand to make the movement, this triggered the button box to record movement onset. Participants reached under the shutter glass to perform the action, as monitored by the experimenter. They were instructed at the beginning of each block of trials which action to perform for the duration of the block: clockwise grasp and lift, counter-clockwise grasp and lift, clockwise reach and knuckle, or counter-clockwise reach and knuckle (Fig. 1C). For grasping trials, participants were instructed to grasp the object using thumb and index finger in the correct orientation, then place the object to the left of the mount for the experimenter to retrieve. For knuckling trials, participants were instructed to tap the centre of the object with the knuckle of their index finger. The end of the movement time was defined as the time when participants’ hand crossed a curtain of infrared beams mounted between two 15 cm tall pillars that were positioned 40 cm apart, immediately in front of the object. Finally, participants returned their hand to the start position on the button box. The experimenter manually marked invalid trials (i.e., incorrect actions or dropped objects).

Each block contained 56 trials (2 shapes x 2 materials x 14 repetitions in random order) during which participants always performed the same action (e.g., clockwise grasp and lift). Actions were randomly assigned across a total of 24 blocks that were completed across 2 sessions of about 3 hours each, over two different days. Breaks were provided in between blocks.

### EEG Acquisition and Preprocessing

EEG data were recorded using a 64-electrode BioSemi ActiveTwo recording system (BioSemi B.V., Amsterdam, Netherlands), digitized at a rate of 512 Hz with 24-bit A/D conversion. The electrodes were arranged in the International 10/20 System. The electrode offset was kept below 40 mV.

EEG preprocessing was performed offline in MATLAB using EEGLAB Toolbox (Delorme and Makeig, 2004) and ERPLAB Toolbox (Lopez-Calderon and Luck, 2014). First, signals were bandpass filtered (noncausal, 2^nd^ order Butterworth impulse response function, 12 dB/oct roll-off) with half-amplitude cut-offs at 0.1 and 40 Hz. Noisy electrodes which correlated <0.6 with nearby electrodes were interpolated (2 electrodes per subject on average). All electrodes were then re-referenced to the average of all electrodes. Independent component analysis (ICA) was performed on continuous blocks for each participant to identify and remove components that were associated with blinks (Jung et al., 2000) and eye movements (Chaumon et al., 2015; Drisdelle et al., 2017). The ICA-corrected data were then segmented relative to the onset of Preview (−100 to 500 ms) and Go Signal (Go; −100 to 500 ms). In addition, invalid trials and epochs containing irregular reaction times (less than 150 ms or greater than 800 ms) were removed. As a result, an average of 0.8% of trials (range: 0.2% −1.8%) from each subject were removed from further analyses.

### Pattern Classification of ERP Signals Across Time

To improve the signal-to-noise ratio of spatiotemporal patterns, we averaged epochs into ERP traces (Grootswagers et al., 2017). First, all blocks of a specific action were pooled together (e.g., clockwise grasp). Up to 14 epochs within a given block that corresponded to the same object (e.g., steel flower) were averaged. This procedure resulted in 6 separate ERP traces per condition (e.g., clockwise grasping of steel flowers) for Preview and for Movement Onset, respectively. Next, we performed multivariate noise normalization by calculating a covariance matrix for all time points of an epoch separately within each condition, then averaging these covariance matrices across time points and conditions (Guggenmos, Sterzer, & Cichy, 2018). We then *z*-scored traces across time and electrodes, and outliers were thresholded at ±3 SD from the mean (for similar approaches: Nemrodov et al., 2017; 2018). We chose this winsorization approach to curb the impact of outliers on SVM-based pattern classification while keeping the number of features constant.

Further, ERP traces were divided into temporal windows with 5 consecutive bins (5 bins * 1.95 ms ≈ 10 ms) to increase the robustness of pattern analyses. For each time bin, data from all 64 electrodes were concatenated to create 320 features. These features were to allow for pattern classification across time, window by window.

We conducted pairwise discrimination of Shape, Material, (hand) Orientation and Action using linear support vector machines (SVM; c = 1; LibSVM 3.22, Chang and Lin, 2011) and leave-one-out cross-validation. That is, for Orientation and Action, 47 of 48 pairs of observations were used for training and one pair was used for testing. For Shape and Material, 23 of 24 pairs of observations were used for training while 1 pair was used for testing. These analyses, except for Action classification, were performed separately on grasping and knuckling data (Action classifiers differentiated grasping vs. knuckling classes). Further, observations for Orientation and Action analyses were drawn from blocks in a random fashion (i.e., same number of data points to create an ERP, but from different blocks) to account for blocked effects (because the same action and hand orientation was performed within the same block).

We also conducted pairwise discrimination for integrated representations of task features, namely, Shape ∩ Material, Shape ∩ Orientation, and Material ∩ Orientation separately for grasping and for knuckling (we did not examine Shape ∩ Material ∩ Orientation representations due to the lack of statistical power). For each of these integrated representations, we always trained four SVMs to reduce the chance that they learned to classify an object based on a single feature alone (although the strategy did not eliminate that possibility completely as explained later). For example, to map Shape ∩ Material representations for grasping we only used ERPs from grasping trials to train a first SVM classifier to distinguish “steel pillows” (class 1) from “steel flowers” and “wood pillows” (class 2) so that shape and material together yielded better classification than either feature alone. A second classifier learned to distinguish “wood pillows” (class 1) from “wood flowers” and “steel pillows” (class 2), a third classifier learned to distinguish “steel flowers” (class 1) from “steel pillows” and “wood flowers” (class 2), and a fourth classifier learned to distinguish “wood flowers” (class 1) from “wood pillows” and “steel flowers” (class 2). Because of the unbalanced nature of the number of observations in each class, we set the cost parameter of the minority class 1 to be double the weight of the majority class 2 (c = 2 vs. c = 1; Batuwita & Palade, 2013) to avoid that the classifier skewed towards the majority class. Next, we performed leave-one-out cross-validation, such that although training data had an unequal number of instances for each class, testing had one of each class (33 observations used for training and 2 for testing, where half of the testing data were from majority class 1 and half were from majority class 2) to further discourage the classifier from using only one of the two features, shape or material. We then averaged the classification performance of the four classifiers to obtain an estimate of the integrated Shape ∩ Material representations for grasping. Likewise, we determined Shape ∩ Material representations for knuckling, and we determined Shape ∩ Orientation and Material ∩ Orientation representations for grasping and knuckling, respectively. For all analyses, decoding was performed for each subject separately and then averaged across participants. Finally, we tested whether integrated classifiers identified truly integrated representations. If so, an integrated classifier should outperform two concurrent classifiers that each used only on a single feature (e.g., only shape and only material, respectively, based on different subsets of electrodes). To this end we first calculated how a single-feature classifier would perform in our test:

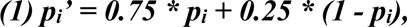

where ***p_i_*** is the probability of some single-feature classifier ***i*** and ***p_i_’*** is its performance in our test (e.g., a shape classifier that attains an accuracy of, say, p_shape_=0.7 would classify a minority object “steel pillow” and a majority object “steel flower” as belonging to class 1 and 2, respectively, with p=0.7 each, and it would classify the second majority object “wood pillow” with p=1-0.7 as belonging to class 2. I.e., considering how often the three objects are tested, this would amount to correct classification with p_shape_’ = 0.5 * 0.7 + 0.25 * 0.7 + 0.25 * (1-0.7) = 0.75 * p_shape_ + 0.25 * (1-p_shape_) = 0.6). Next, to estimate how two single-feature classifiers perform optimally together we modelled the response of each single-feature classifier as a continuous value (i.e., as the signed distance from the decision boundary) with a Gaussian distribution where ***µ*** is the mean, ***σ_i_*** is the standard deviation, and ***p_i_’*** is the area under the Gaussian up to a decision boundary ***x***. The p_i_’ value can then be written as a cumulative Gaussian:

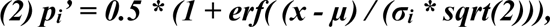

where ***erf*** is the error function and ***sqrt*** is the square-root. Solving equation (2) for σ_i_ for both single-feature classifiers (and with arbitrary values for µ and x), allows us to apply maximum likelihood estimation to calculate the optimally combined standard deviation for both single-feature classifiers together:

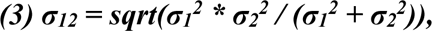

where ***σ_1_*** and ***σ_2_*** are the standard deviations for the two single-feature classifiers, respectively, and ***σ_12_*** is the standard deviation of the optimally combined classifier. Inserting ***σ_12_*** into equation (2) gives us the optimal performance of both classifiers together (one exception: if one or two classifiers performed at guessing rate or worse, optimal performance p_12_ was set to the maximum of p_1_, p_2_, or 0.5). Hence, the accuracy of the integrated classifier was tested relative to p_12_.

### EMG Acquisition and Analysis

We recorded surface EMG from four muscles (superior trapezius, anterior deltoid, brachioradialis, and first dorsal interosseous) at 2000 Hz using a BioSemi ActiveTwo recording system. Signals were rectified and then high-pass filtered (noncausal Butterworth impulse response function, fourth order) with a cutoff frequency of 2 Hz. Subsequently, we computed a root mean square (RMS) value of the signal at each point in time using a 10 ms gliding window. Signals were then down-sampled to 512 Hz and segmented relative to the onset of Preview or to movement onset.

We approached decoding time-resolved EMG patterns with the same logic as the EEG data (see Pattern classification of ERP signals across time; Figure 6).

### Cross-Temporal Generalization of ERPs

Temporal classification examined whether classifiers trained on EEG data from a specific time point is capable of decoding data from the same time point. However, for cross-temporal generalization we extended this approach, whereby we examined whether classifiers trained from a certain time can decode data from the same or different time points. That is, we trained and tested classifiers on any combination of times from Preview to be plotted in a square-shaped graph with two time axes (e.g., Fig. 3). To understand differences and similarities in the representation of features modulated by action, we conducted separate analyses: training and testing on grasping, training and testing knuckling, and the average of training on grasping and testing on knuckling and vice versa. If the underlying representations are similar between two selected time windows, then it is expected that a classifier will generalize between these time windows. Specifically, we expected to see three patterns of classifier performance reflecting different neural representations: 1) Chained representations. The classifier will perform above chance only when trained and tested on the same time points, yielding a classification pattern along the diagonal of the graph. This pattern indicates that classification is driven by a chain of transient neural representations that activate sequentially. 2) Reactivated representations. The classifier trained at an earlier time will perform above chance for later time points. This results in significant classification along the diagonal as well as at separate and symmetrical areas above and below the diagonal and reflects that later neural representations reactivate earlier ones. 3) Sustained representations. The classifier trained at an earlier time will perform above chance when tested for times immediately following the training time, resulting in a wide coverage of areas above and below the diagonal. This suggests that representations are sustained across time.

### Pattern Classification of Time Frequency Data

To perform classification on time frequency transformed data, epochs were first bandpass filtered (delta: 1-3 Hz; theta: 3-8 Hz; alpha: 8-12 Hz; beta: 15-25 Hz; gamma: 30-40 Hz) using two-way least-squares finite impulse response filtering. We then applied Hilbert transform on narrowband data to obtain the discrete-time analytical signal and squared complex values to attain the power. This band-specific power was otherwise pooled, normalized and averaged in the same way as ERP traces for spatiotemporal EEG classification (see “Pattern Classification of ERP Signals Across Time” and “Cross-Temporal Generalization of ERPs”).

### Electrode Informativeness

To assess the informativeness of electrodes for the classification of ERP effects (Shape, Material, Orientation, and Action) we conducted a searchlight analysis across electrodes. For each electrode we defined a 50 mm radius neighbourhood and performed classification on spatiotemporal features obtained from each electrode neighbourhood across 100 ms time bins.

### Significance Testing

For behavioural data we determined statistical differences (reaction and movement time) using four-way repeated measures ANOVAs (Action x Hand Orientation x Shape x Material), adjusting for sphericity violations using Greenhouse-Geisser (GHG) corrections. Additional post hoc analyses were conducted using repeated measures t-tests and corrected for multiple comparisons using the false discovery rate (FDR; Benjamini & Hochberg, 1995).

For all tests conducted on EEG data we assessed statistical significance using a non-parametric, cluster-based approach to determine clusters of timepoints (or electrodes for searchlight analyses) where there were significant effects at the group level (Nichols and Holmes, 2002). In the case of time-resolved analyses, we defined clusters as consecutive time points that exceeded a statistical threshold. Specifically, cluster thresholds were defined by the 95^th^ percentile of the distribution of t-values at each time point attained using sign-permutation tests computed 10000 times, equivalent to p<0.05, one-tailed. Significant clusters were determined by cluster sizes equivalent to or exceeding the 95^th^ percentile of maximum cluster sizes across all permutations (equivalent to p<0.05, one-tailed). Similarly for searchlight analyses carried out across electrodes, cluster-based correction was performed on each 100 ms time window, separately on spatial clusters of electrodes.

For temporal generalization analyses, we tested for significant differences between three types of representations (i.e., chained, reactivated, and sustained representations) during Preview before grasping vs. knuckling (Fig. 3). To this end we calibrated regions of interest (ROIs) based on the pattern observed for shape representations during grasping, because this “grasp-shape” condition showed clear reactivated representations that allowed us to define distinct regions indicative of all three types of representations. 1) To create a *diagonal* ROI we manually selected the set of all significant points clustered along the main diagonal of the graph for the grasp-shape condition (based on visual inspection, an area that ranged from approximately 110 ms to 500 ms on the x and y axes). This resulted in 277 points to be included in the diagonal cluster. Next, we calculated the horizontal and vertical centre of the cluster, and we determined the cluster’s standard deviations both parallel and orthogonal to the diagonal across participants. Lastly, we used these measures to define a diagonal rectangle centred on the diagonal cluster with sides that were +/-1.7 times its standard deviations. We chose a factor of 1.7 such that the ROIs would cover the largest area of significant points without overlapping with the next two ROIs.

2) Next, we defined an *“arms-shaped”* (i.e. “arms” off of the diagonal) ROI: we selected the significant cluster below the diagonal found in the grasp-shape result (i.e., the off-diagonal cluster that was larger). Specifically, we selected 72 significant points below the diagonal corresponding to significant areas in the time range ∼250-500 ms on the x-axis and ∼100-200 ms on the y-axis. Given this cluster, we set up a rectangle with the same horizontal and vertical mean and with vertices that were +/-1.7 times the cluster’s horizontal and vertical standard deviations. We then inverted the x and y coordinates of the rectangle to form the second “arm”, in the symmetrical above-diagonal area corresponding to the approximate time range of 100-200 ms on the x-axis and 250-500 ms on the y-axis.

3) To define a *“triangular”* ROI for *sustained* representations, we selected triangular areas abutting the chained and reactivated ROIs. Specifically, the triangles had a base width the same as the length of the respective “arm”, and with the height defined by the end coordinate of the diagonal ROI.

4) In addition, we defined a baseline ROI that encompassed all data points with training or testing times smaller than 50 ms, i.e., for times where classification should be at or near chance.

For each of these ROIs we calculated, for grasping and knuckling separately, a difference score of the average grasping vs. knuckling accuracies. Next, we determined one-tailed 95% confidence intervals (because a priori we expected representations during grasping, if at all, to be more prominent than representations during knuckling) using non-parametric bootstrapping with 10000 iterations, and considered a grasping vs. knuckling difference for the diagonal, “arms-shaped,” or triangular ROI to be significant if it was more positive than the difference for the baseline ROI with no overlap in confidence intervals. We repeated this significance testing process for all main effect and interaction comparisons between grasping and knuckling such that a significant difference within the diagonal ROI would signify a stronger chained representation, a difference within the arms-shaped ROI (but not the triangular ROI) would indicate stronger reactivated representations, and better decoding within the triangular ROI (with or without differences in the arm-shaped ROI) would indicate a stronger sustained representation.

## Results

### Behavioural Results

The average reaction time (RT; time between Go onset and movement onset) was 307 ms (SD = 78 ms). RTs submitted to a four-way repeated-measures ANOVA (Action x Hand Orientation x Shape x Material) yielded a main effect of Hand Orientation (F (1, 15) = 4.878, p < 0.05 η_p^2 = 0.245) such that reaction times were longer for counter-clockwise hand orientations (mean RT = 320ms) than clockwise hand orientations (mean RT = 295ms). No other main effect or interaction was significant (F’s < 3.708, p’s > 0.073).

The average movement time (MT; time between movement onset and movement end) was 249 ms (SD = 32 ms). The four-way repeated-measures ANOVA of MTs showed a main effect of Action (F (1,15) = 33.634, p < 0.001, η_p^2 = 0.692), such that knuckling (mean MT = 279ms) was slower than grasping (mean MT = 220ms). All other effects were non-significant (F’s < 3.565, p’s > 0.079).

### Classification of ERPs for Main Effects

Given our main objective we looked at classification for Shape and Material (as well as for Hand Orientation) separately for grasping and knuckling. Classification accuracy for Grasping Shapes representations (Fig. 2A, first row, dark blue) first became significant briefly from 70 ms to 80 ms. From 100 ms onwards, accuracy continued to rise and peaked at 170 ms, then becoming stable until the end of the Preview period. Classification accuracy for Knuckling Shapes became significant later than grasping, from 140 ms to 430 ms (interruptions at 250-270 ms and 320-340 ms). There was only a transient period where Grasping Shapes classification accuracy was significantly higher than Knuckling Shapes accuracies (from 100 ms to 120 ms). Material classification for grasping was significant during three periods: 140-170-ms, 240-280 ms, and 300-320 ms (Fig. 2A, second row, dark red). For knuckling, the material curve rose earlier, becoming significant non-continuously from 140-320 ms (140-250 ms; 270-320 ms), and later, from 340-430 ms. However, we observed no difference in material classification for grasping vs. knuckling. Orientation classification was robust for both grasping (starting at 50 ms; Fig. 2A, third row, dark green) and knuckling (starting at 70 ms). Both curves rose quickly and remained significant for the duration of the Preview period. There was a short time from 300 to 330 ms where grasping classification of orientation was higher than knuckling. Lastly, action classification became significant early at 60 ms (Fig. 2A, fourth row) and remained robustly significant throughout Preview.

**Figure 2.**
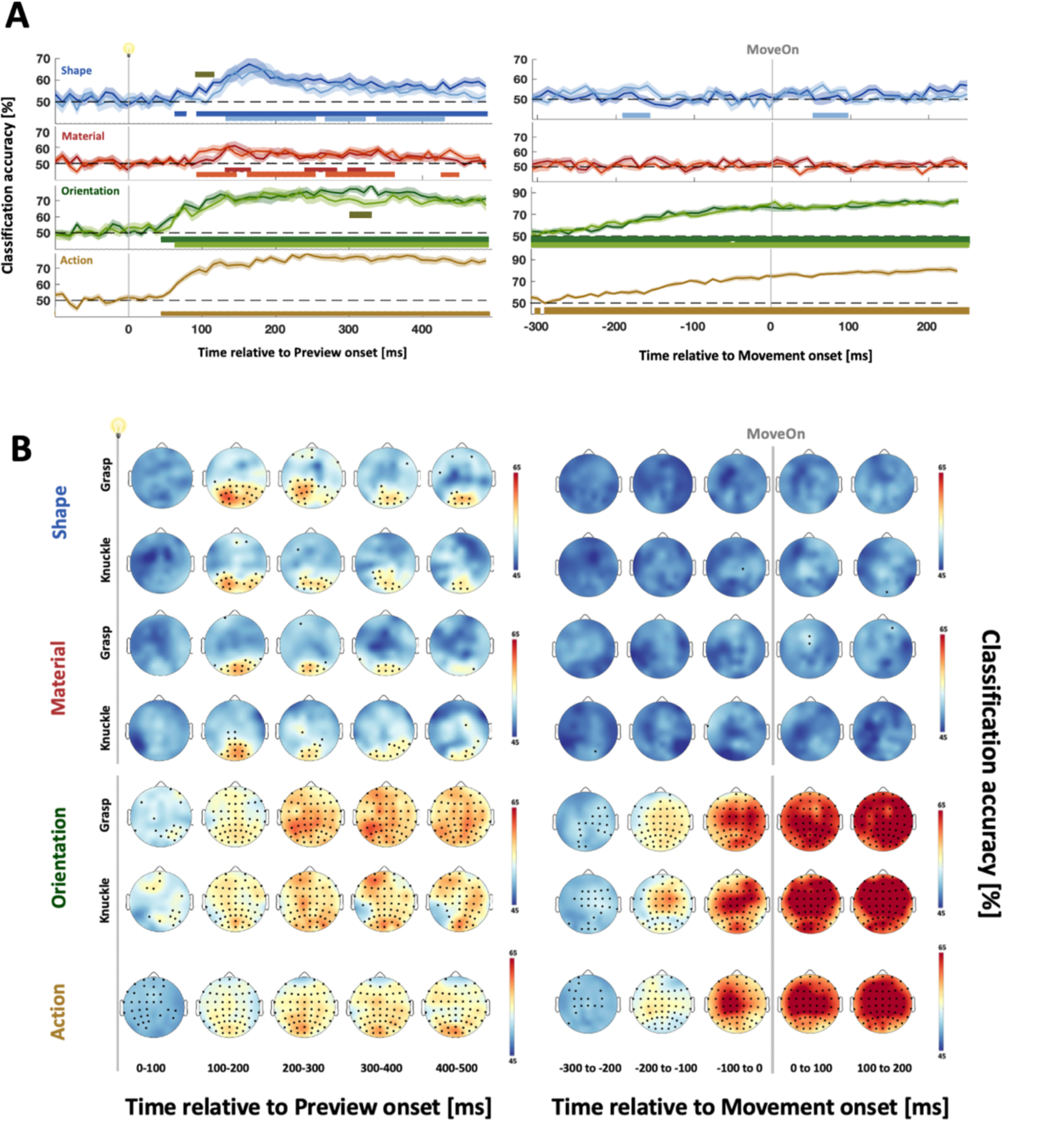
(A) Time resolved classification of main effects aligned to Preview (left) and Movement Onset (right). Darker curves represent grasping, lighter represent knuckling. Horizontal coloured bars denote significantly above chance classification (cluster-corrected t test, one-tailed; p<0.05). Grey horizontal bars denote significantly better classification for grasping compared to knuckling (cluster-corrected t test, one-tailed; p<0.05). (B) Electrode informativeness for respective main effects aligned to Preview (left) and Movement Onset (right; two-tailed one-sample t test; q < 0.05). MoveOn: Movement Onset.

For data aligned to Movement Onset, classification for Shape and Material did not reach significance (with an exception for Knuckling Shape from −210 ms to −240 and 50 to 100 ms, see Fig. 2A first row, right side). Classification for Hand Orientation was significant prior to movement (average RT 307 ms), peaking around movement onset and remaining high for the rest of the movement time (exception a brief interruption at 50 ms prior to Movement Onset for grasping classification). Action classification followed a similar trajectory to orientation (becoming significant before movement onset, peaking at movement onset and remaining at a high level for the rest of movement time).

### Electrode Informativeness

Shape during grasping was most strongly represented in electrodes concentrated over left occipital regions (including O1, PO7, PO3, P5 and P3) during the first 200 ms after stimulus presentation (Fig. 2B, first row). Representations continued to be more lateralized to the left hemisphere from 300-400 ms, the strongest informativeness moving more central (including P5, P3, P1, CP3 and CP1). From 300 ms onwards, posterior occipital electrodes showed the greatest informativeness (namely, PO3, POz and PO4). Similar to grasping, shape representations during knuckling were initially more strongly lateralized to the left (100-200 ms, including electrodes O1, PO7, PO3, P5 and P3), thereafter being more concentrated on posterior occipital electrodes over both hemispheres (including PO3, POz, Oz and O2; exception at 300-400 ms where posterior central electrodes were also informative: CP3, CP1, CP2). Material representations in contrast, did not show lateralization in either grasping or knuckling data. Occipital electrodes most greatly contributed to classification, especially from 100-200 ms (Fig. 2B, third row, grasping: Oz, O2, POz, P4, P8; knuckling: O1, Oz, O2, PO3, POz, PO4). Orientation (Fig. 2B, fifth and sixth rows, both grasping and knuckling), as well as Action (Fig. 2B, seventh row) showed an involvement of widely distributed electrodes during Preview.

For data aligned to movement onset, Shape and Material yielded close to no reliable results for electrode informativeness (Fig. 2B, first two rows, right). However, Orientation classification drew information from central electrodes (seen more prominently in knuckling data, including electrodes: Cz, C2, FC1, FCz, FC2) starting 200 ms before Movement Onset and becoming widely distributed 100 ms before Movement Onset and throughout the movement period. As expected, Action classification was most prominently informed by central electrodes over the left hemisphere (most strongly 100 ms before Movement Onset, electrodes: CP3, CP1, CPz, C3, C1, Cz, FC3, FC1 and FCz) before becoming widely distributed after movement onset.

### Temporal Generalization of ERPs

Because we found little evidence for grasp-specific task features influencing representations during grasping vs. knuckling (i.e., a fleeting difference for Shape classification and no difference for Material classification), we performed several additional analyses: temporal generalization, time-frequency analysis, analysis integrated representations of shape and material, and of representations during later phases of action execution. First, we explored the degree to which representations generalized over time. We found that temporal generalization for Grasping Shapes showed a distinct pattern of reactivation, such that times trained at ∼100-200 ms generalized to times tested from 240 ms to 500 ms during Preview and vice versa (Fig. 3A, first graph). This shows that when participants intended to grasp objects, earlier shape representations reactivated at a later time (e.g., King & Dehaene, 2014).

**Figure 3.**
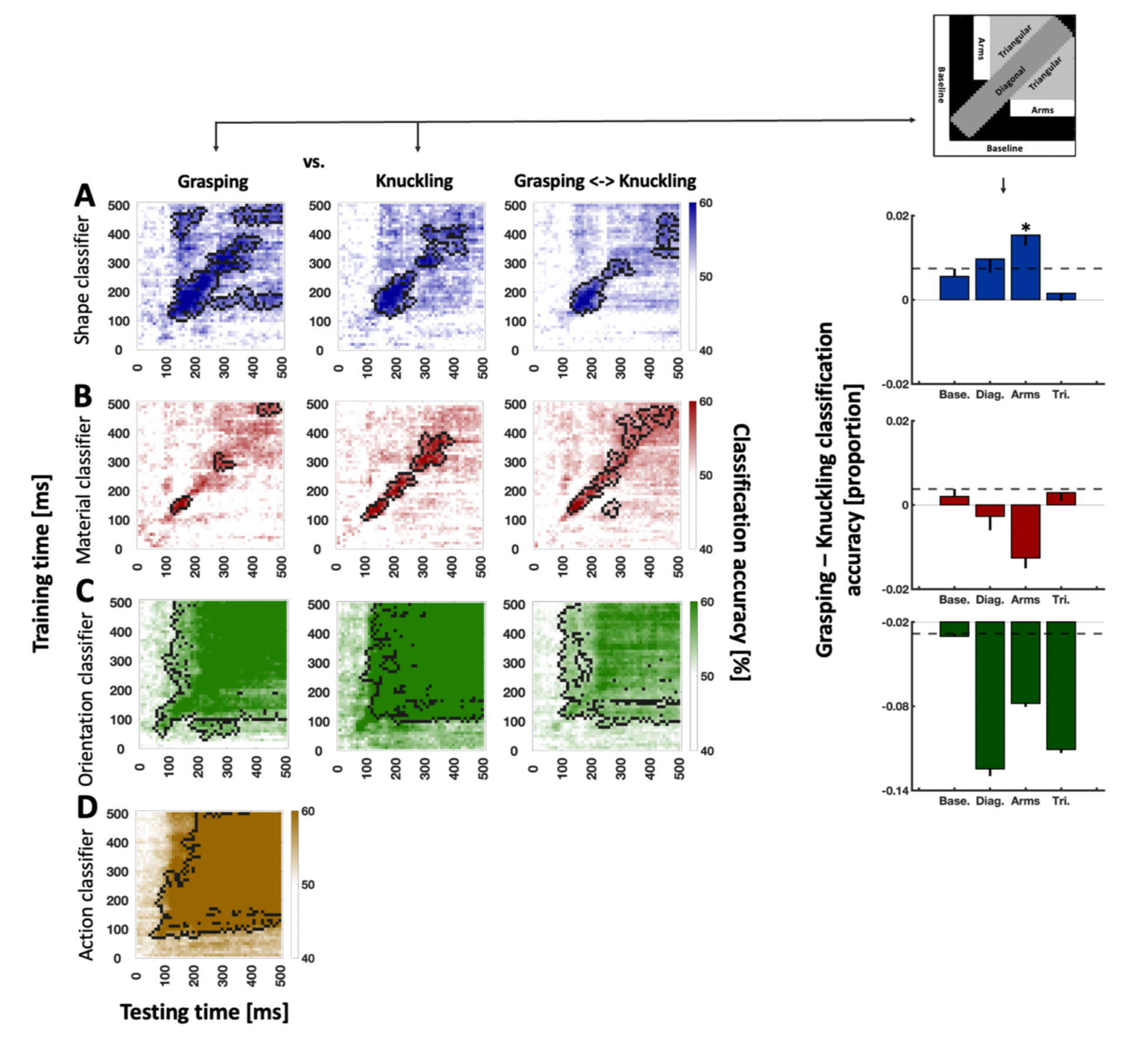
Temporal generalization of Shape (A), Material (B), (hand) Orientation (C), and Action classification (D). Left side of figure: Time-by-time plots for grasping (first plot), knuckling (second plot), and the average of results for grasping trained on knuckling and vice versa (third plot). Note that the plot for action shows classification accuracy for grasping vs. knuckling rather than for grasping and knuckling separately. For all time-by-time plots black outlines indicate significant clusters (cluster-based sign-permutation test with cluster-defining and cluster-size thresholds of p < 0.05). Right side of figure: Top plot depicts ROIs inspected for classification differences between grasping and knuckling. Bar graphs depict observed differences between grasping and knuckling for the ROIs. Base: baseline ROI, Diag: diagonal ROI, Arms: “arm-shaped” ROI, Tri: Triangular ROI. Error bars represent bootstrapped confidence intervals (10,000 iterations, one-tailed). Dashed lines visualize significance level (i.e., the upper boundary of the confidence interval for the baseline ROI). Asterisk represents significant results relative to baseline ROI.

Interestingly however, shape representations did not reactivate for knuckling, i.e., significant classification was confined to the diagonal (Fig. 3A, second graph), suggesting that during knuckling shape representations continuously changed over time. Statistically comparing the different regions of interest (ROIs) for grasping vs. knuckling showed that Grasping Shapes classifiers were significantly more accurate along the “arms” of reactivation whereas there was no difference for the triangular ROI encapsulated by both (Fig. 3A, fourth graph). For the diagonal ROI we observed a trend in that the difference was greater than zero, but this might have been due to unspecific fluctuations between blocks of trials where participants grasped vs. knuckled the objects given the difference for the baseline ROI (dashed line in Fig. 3A, fourth graph). Finally, training classifiers on grasping data and testing on knuckling data (and vice versa) showed that shape classification along the diagonal was comparable for grasping and knuckling (Fig. 3A, third graph). In sum, we found that similar shape representations formed regardless of action intention but that earlier representations (i.e., similar brain areas or networks of areas) reactivated when there was a specific need for more detailed shape processing during grasping.

Next, we looked for temporal generalization of Material representations but only found significant classification along the diagonal (for grasping, knuckling, and for the generalization between grasping and knuckling) (Fig. 3B). Also, when comparing ROIs we found no evidence for our a-priori assumption that material representations should be more prominent when material is task-relevant (i.e., during grasping). By contrast, there was a trend in the opposite direction, especially for the “arms-shaped” ROI (Fig. 3B, fourth graph) which was not captured by the one-tailed testing approach of the current study. It is possible that the trend reflects that the intention to knuckle the objects directs a focus of attention to the surface (i.e., the texture) of the objects and thereby amplifies representations of material. However, more research is required to explore this possibility.

We also inspected temporal generalization for Orientation observing robust classification along the diagonal and in the regions above and below the diagonal. This pattern is indicative of sustained representations during grasping and knuckling (Fig. 3C, first and second graph; n.b., classification during knuckling was numerically more accurate than during grasping, Fig. 3C, fourth graph, but once again that difference was not captured by our a priori interest in representations that are more prominent during grasping and, thus, will require further testing in the future). Interestingly, training classifiers on grasping data and testing on knuckling (and vice versa) yielded a similarly robust pattern of temporal generalization, suggesting that Hand Orientation representations were mostly similar regardless of action intention (Fig. 3A, third graph).

Lastly, we examined temporal generalization for Action. We found that Action representations, similar to Hand Orientation representations, generalized above and below the diagonal, suggesting that Action representations attained a sustained state over time (Fig. 3D).

### Time Frequency

To explore whether the grasp-specific reactivation of Shape representations reflected attentional mechanisms, we examined individual frequency bands where alpha and theta band have been previously shown to change with regards to spatial and object-based attention, respectively (Harris et al., 2017; Liebe et al., 2012; Capotosto et al., 2009). Indeed, we found that classification of Shape yielded significant results that overlapped in time with the Shape effects (Fig. 2A and 3A) in both frequency bands (alpha for Knuckling Shapes: 170 – 240 ms; theta for Grasping Shapes: 110 – 300 ms; theta for Knuckling Shapes: 90 – 310 ms; no reliable difference between Grasping and Knuckling Shapes). To explore these effects further we conducted a temporal generalization analysis for alpha and theta band. Neither of the two bands yielded significant results compared to chance (Figure 4B, left side). However, for the pre-defined ROIs we found that Shape classification was more accurate for grasping than knuckling along the diagonal compared to the baseline ROI in both frequency bands and there was a non-significant trend for the “arms-shaped” ROI within the alpha band (Fig. 4B, right side).

**Figure 4.**
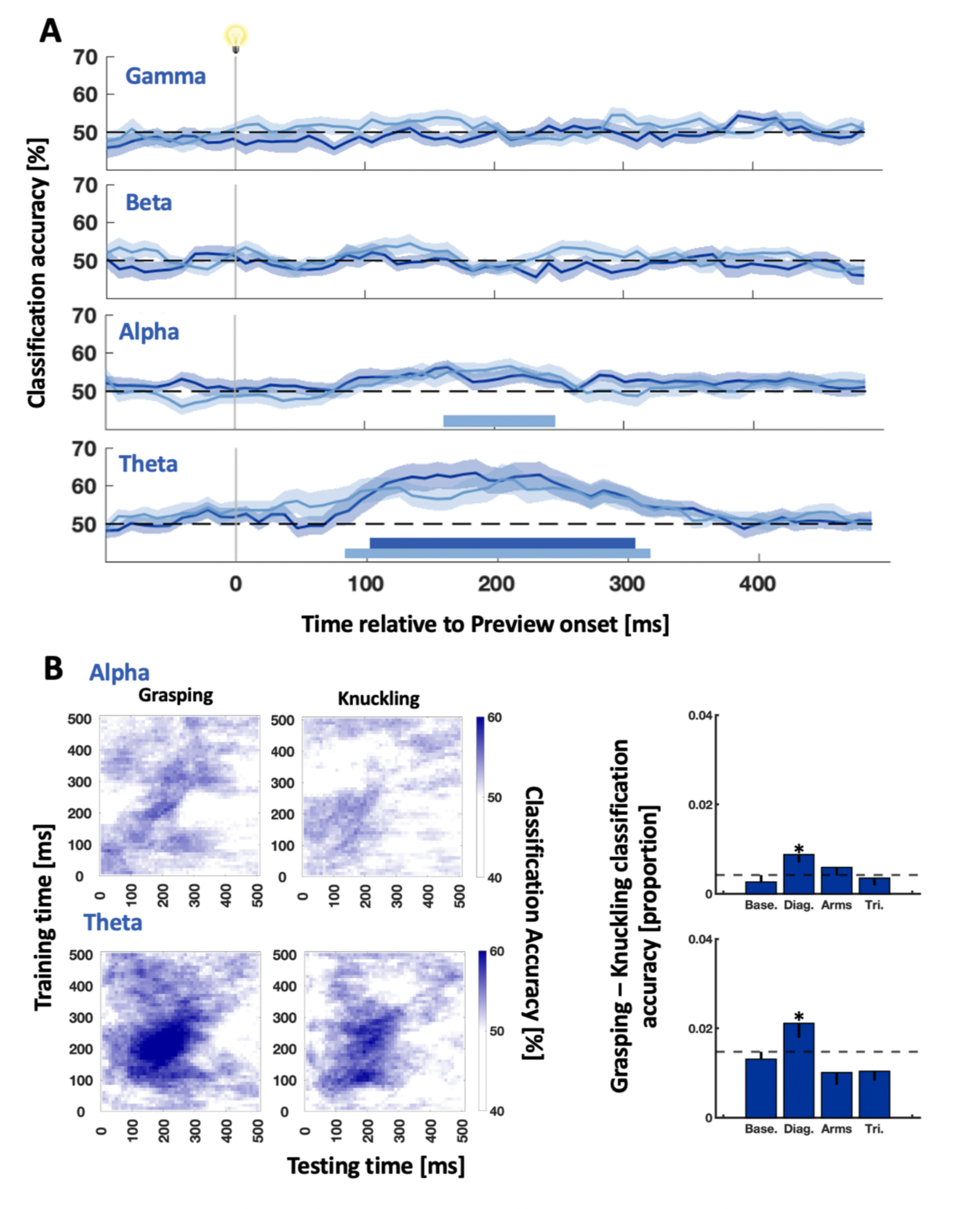
(A) Time course of Shape classification for time frequency data for gamma, beta, alpha and theta bands. Darker blue: grasping data, lighter blue: knuckling data. Horizontal coloured bars represent significantly above chance classification accuracies (cluster-corrected t-test, one-tailed; p<0.05). (B) Temporal generalization of shape classification for alpha (first row) and theta band (second row). Left side of (B): Time-by-time plots for grasping (first plot) and knuckling (second plot) (classification not significantly different from chance). Right side of (B): bar graphs depict differences between grasping and knuckling for the pre-defined ROIs based on Fig. 3. Base: baseline ROI, Diag: diagonal ROI, Arms: “arm-shaped” ROI, Tri: Triangular ROI. Error bars represent bootstrapped confidence intervals (10,000 iterations, one-tailed). Horizontal dashed lines visualize significance level (i.e., the upper boundary of the confidence interval for the baseline ROI). Asterisks represent significant results relative to baseline ROI.

### Integrated Representations of Task Features

Next, we tested whether grasp-specific representations formed in an integrated manner, e.g., as representations for a specific shape joined together with a specific material or orientation, or a specific material joined with a specific orientation. Only the Shape ∩ Orientation grasping classifier performed better than that for knuckling (110-130 ms and 230-260 ms, Fig. 5, second row). That said, none of the integrated classifiers performed better than predicted by the respective two single-feature classifiers (e.g., Shape ∩ Material classifier vs. shape classifier and material classifier etc.).

**Figure 5.**
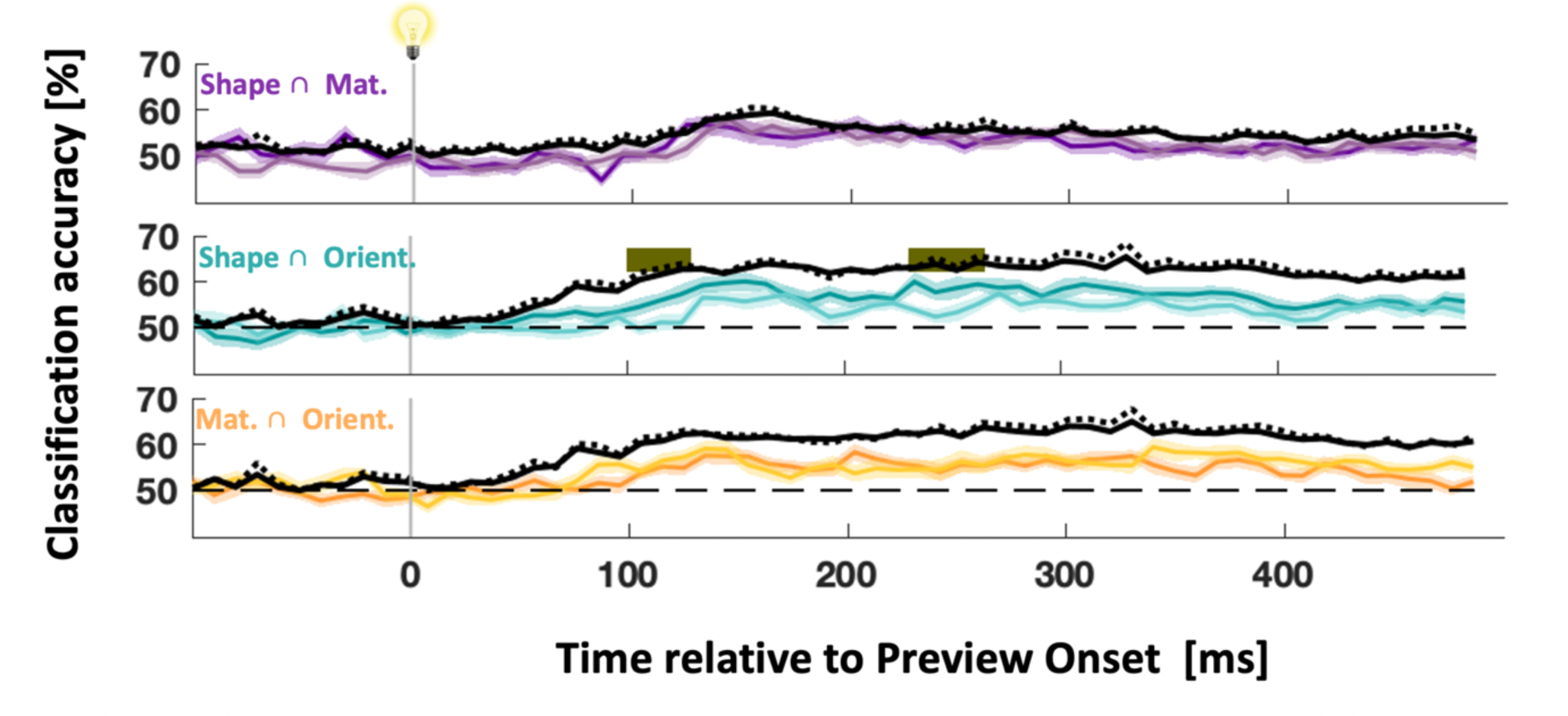
Time course of classification for interactions between Shape ∩ Material (purple), Shape ∩ Orientation (turquoise) and Material ∩ Orientation (orange). Darker coloured curves show classification for grasping, lighter curves for knuckling. Black curves show performance as predicted by the respective two single-feature classifiers for grasping (solid line) and knuckling (dotted line). Grey horizontal bars denote significantly better classification for grasping than knuckling (cluster-corrected t test, one-tailed; p<0.05).

### ERP and EMG Classification Broken down by Movement Phase

Because all previous analyses produced no evidence for grasp-specific Material representations during Preview, we reasoned that the material of the objects should become relevant at least when participants were about to lift the objects. Indeed, we found Material representations and their integrated representations with Shape and Orientation to perform significantly better for grasping than knuckling data; there were grasp-specific difference for Material after object contact: first approximately during the load phase (400– 440 ms) and then again during the lift phase (650 – 830 ms, Fig. 6A, second row). Importantly, at least the effect during load was unlikely to be contaminated by muscle activity because it preceded grasp-specific classification based on EMG data (starting at 440 ms Fig. 6B, second row). Next, integrated representations of Shape ∩ Material showed a trend to be stronger for grasping than knuckling from −20 to 40 ms, i.e., well before grasp-specific Shape ∩ Material classification of EMG data (Fig. 6A, third row vs. Fig. 6B, third row; a later grasp-specific Shape ∩ Material representation, 840 – 900 ms, occurred after the respective EMG classification became significant). That said, performance was not better than predicted by the single-feature classifiers for shape and material. Material ∩ Orientation representations were also more prominent for grasping than knuckling (410 – 460 ms and non-continuously from 540 – 890 ms, Fig. 6A, fifth row) and, thus, became significant shortly before grasp-specific classification of EMGs (beginning at 510 ms; Fig. 6B, fifth row). But once again, performance was worse than that predicted by the single-feature classifiers for material and orientation.

**Figure 6.**
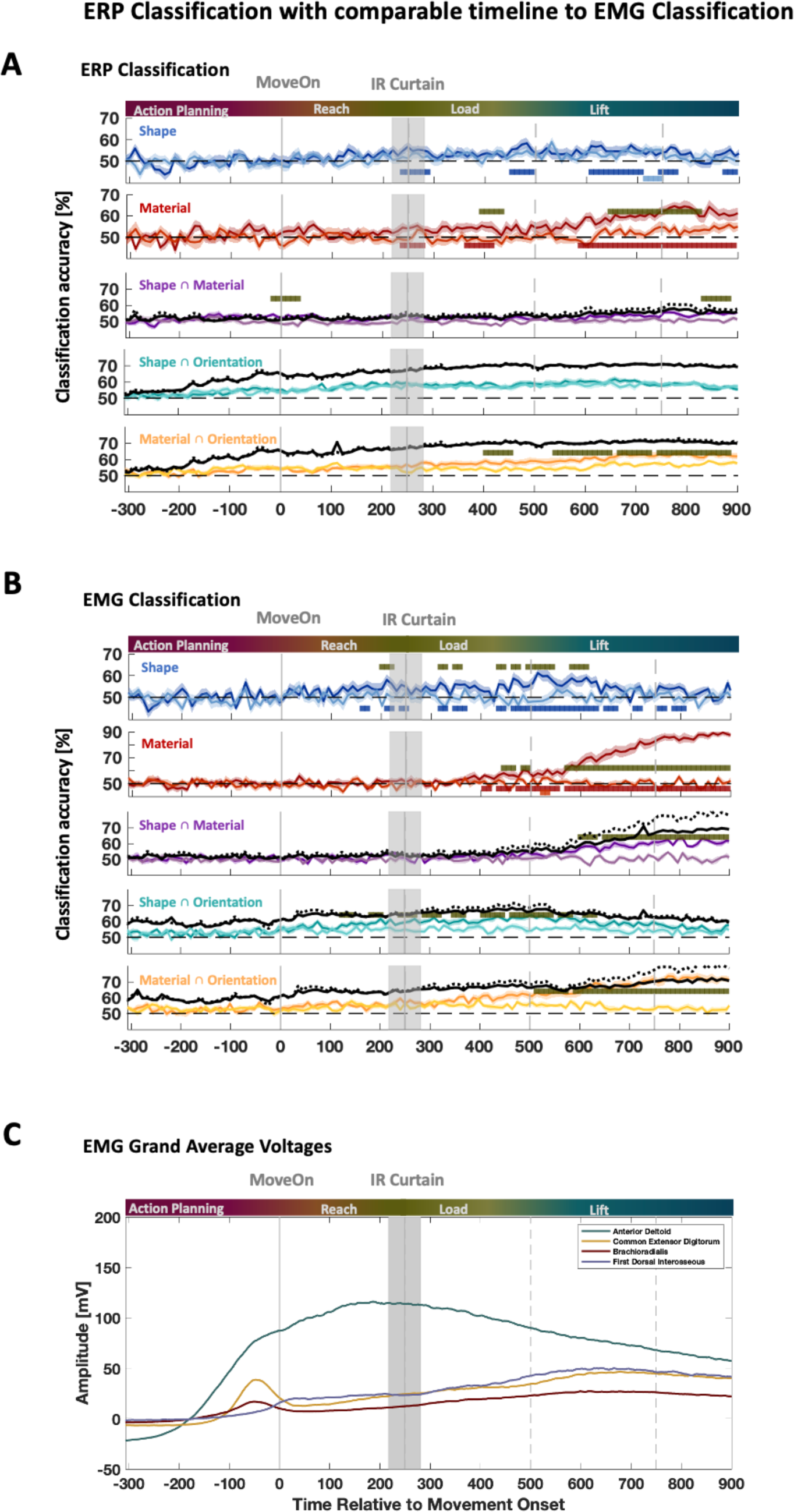
ERP and EMG data aligned to Movement Onset. (A) Time-resolved classification of ERPs for Shape, Material, Shape ∩ Material, Shape ∩ Orientation and Material ∩ Orientation (B) Time-resolved classification of EMGs. (C) EMG grand average voltages for recorded muscles as they pertain to movement phases. (A&B) Darker coloured curves show grasping data, lighter curves are knuckling data. Black curves show performance as predicted by the respective two single-feature classifiers for grasping (solid line) and knuckling (dotted line). Coloured horizontal bars denote significantly above chance classification accuracies (cluster-corrected t test, one-tailed; p<0.05). Grey horizonal bars denote significantly better classification accuracies for grasping compared to knuckling (cluster-corrected t test, one-tailed; p<0.05). (A-C) Multi-coloured bands at the top of each sub-plot indicate approximate movement phases based on grand average ERPs (Fig.6C). MoveOn: Movement Onset. IR Curtain: time at which participants’ hand intersected the infra-red curtain mounted immediately in front of the objects.

## Discussion

In the present study we tested whether the brain forms action-specific representations of task features during grasping as a well-rehearsed sensorimotor task. Specifically, we recorded EEG from the scalp of participants while they either grasped objects or touched them with the knuckle of their index finger, and we used objects that differed in shape and material (i.e., weight), hence, features that are relevant for grasping but not for knuckling. Utilizing the sensitivity of SVMs to analyze our EEG data, we found evidence that, specifically for grasping, the brain formed representations of shape during movement planning, and representations of material after object contact.

In detail, we found that viewing objects triggered a chain of neural representations of object shape that was largely comparable regardless of whether participants intended to grasp or knuckle objects, suggesting that visual object analysis was mostly the same for both actions. Crucially however, only for grasping did we find that earlier shape representations (from about 100 to 200 ms after object onset) activated again later (from 240 ms to 500 ms).

This reactivation is consistent with a previous EEG study where reactivation occurred at about the same time during a grasp task (Guo et al., 2019). Importantly however, we now show that reactivation is specific for grasping and not knuckling.

This finding resembles, to some extent, reactivation as observed with fMRI where reactivation occurred in a delayed memory-guided grasping task (Singhal et al., 2013; Monaco et al., 2017), although the latter studies examined reactivation on a quite different time scale of several seconds and involved memory functions that were not required for the present paradigm.

Instead, the grasp-specific reactivation as observed here complements another imaging study showing that during object preview early visual areas carry detailed grasp-related information (Gallivan et al., 2019). Together the findings indicate that the intention to grasp an object is associated with sensorimotor control signals that feed back to earlier visual areas, arguably for a more detailed analysis of the object as required for grasp computations, perhaps similar to a database query.

These mechanisms could reflect action-driven forms of visual attention where attentional foci arise near the selected grasp points on an object (Baldauf & Deubel, 2009; Baldauf et al., 2006; Brouwer et al., 2009; Schiegg et al., 2003). Consistent with this idea, we found that shape representations were particularly prominent within alpha and theta band that are sensitive to processes of spatial and feature-based attention (Capotosto et al., 2009; Harris et al., 2017; Liebe et al., 2012). Grasp-specific shape representations, especially within the alpha band, exhibited an activation pattern that was somewhat similar to that found for ERP data, exhibiting a chain of shape representations that was more prominent for grasping than for knuckling and a trend for enhanced reactivation. This indicates that intention-specific attentional mechanisms accompanied preparatory grasp computations.

Unlike grasp-specific representations of shape during movement preparation, no such specific representations were found for material, and that was also true for integrated grasp-specific shape ∩ material representations. This null result for movement planning was not due to poor signal-to-noise ratios, given that material classification was robust for grasping as well as for knuckling. Furthermore, we did find grasp-specific material representations and a trend for grasp-specific shape ∩ material representations, but these effects occurred during movement execution. Specifically, transient grasp-specific shape ∩ material representations occurred around movement onset (with the caveat that the integrated classifiers were outperformed by single-feature classifiers), and grasp-specific material representations emerged during the load phase consistent with behavioural data (Baugh et al., 2012; Buckingham et al., 2009; Johansson & Flanagan, 2009).

Taking these findings together shows that action-specific representations of task features mostly form separately and at disparate times when the respective feature is needed for a certain phase of the movement. That is, shape information is required for computing reach-to-grasp movements towards the selected grasp points on an object and, thus, shape representations need to occur during movement planning. By contrast, movement trajectories are largely unchanged by visual cues about object weight such as filled vs. empty bottles (Sartori et al., 2011) or, for horizontal grasps, the vertical size objects (despite vertical size indicating object weight; Ganel & Goodale, 2003). Instead, expectations about weight have a significant influence on the load phase (e.g., Johansson & Flanagan, 2009).

Given such a pattern of phase-specific computations it should be expected that features that are indeed relevant for the same movement phase are integrated into a joint representation. Consistent with this idea, we did observe a trend for an integrated grasp-specific representation of shape and hand orientation. Presumably the effect would have been larger had our objects not been designed with identical spatial coordinates for their grasp points but with more different ones, thus, more prominently impacting grasp trajectories. Further research is required to test this prediction. Future research should also explore whether integrated task-specific representations arise in prefrontal cortex where the distributed activity of so-called mixed selectivity neurons reflects the integration of multiple task features (Rigotti et al., 2013), or whether integrated representations arise across multiple specialized brain regions that functionally connect with one another (e.g., Lakatos et al., 2007).

As for the present study, integrated grasp-specific representations should have been expected given abundant evidence that for goal-directed actions task features join into “event files” or conjunctive representations (e.g., Hommel, 2019). However, based on this previous evidence such an integrated representation should have formed soon after the object became visible and should have been evident for longer periods of time (e.g., Kikuomoto & Mayr, 2020) whereas the present study produced some evidence for integrated grasp-specific shape and orientation representations during movement planning whereas trends for integration of shape and material occurred late, around the time of movement onset, and all integrated representations were transient.

We argue that our results diverge from previous findings of conjunctive representations because those paradigms relied on arbitrary combinations of task features and afforded actions. For that reason, they required sustained representations in working memory. In contrast, grasping as used in the present study entails little working memory; it is a highly overlearned task with shape and material being features that participants have a lifetime worth of experience with. Thus, our data argue against the universal validity of event files for any kind of goal-directed behaviour.

In conclusion, in the present study we found evidence for mainly distinct representations of individual task features at times when those features were required for the programming of the different phases of grasping: grasp-specific representations of shape occurred during grasp planning indicating that shape information is incorporated into the computations for grasp-point selection and the programming of reach-to-grasp trajectories. On the other hand, grasp-specific representations of material occurred after object contact suggesting that weight information is entered into grasp computations during the load phase. Together these findings suggest a model of grasp control where grasp-specific processing of task features occurs at times when the respective feature becomes relevant for a specific subcomponent of the action. Hence, grasp-specific feature representations arise upon demand.

## Acknowledgement

This research was supported in part by a grant from the Natural Sciences and Engineering Research Council of Canada (NSERC).

